# Enhanced Yield and Gentle Purification of HIV for Cryo-Electron Tomography Analysis of Virion Maturation

**DOI:** 10.1101/2024.12.12.628087

**Authors:** Benjamin Preece, Wiley Peppel, Rodrigo Gallegos, Gillian Yassasi, Gabe Clinger, Nicole Bohn, Broti Adhikary, Luiza Mendonça, David Belnap, Michael Vershinin, Saveez Saffarian

## Abstract

HIV is a lentivirus characterized by the formation of its mature core. Visualization and structural examination of HIV requires purification of virions to high concentrations. The yield and integrity of these virions are crucial for ensuring a uniform representation of all viral particles in subsequent analyses. In this study, we present a method for purification of HIV virions which minimizes forces applied to virions while maximizing the efficiency of collection. This method allows us to capture between 1,000 and 5,000 HIV virions released from individual HEK293 cells after transfection with the NL4.3 HIV backbone, a 10 fold advantage over other methods. We utilized this approach to investigate HIV core formation from several constructs: pNL4-3(RT:D_185_A&D_186_A) with an inactive reverse transcriptase, NL4.3(IN: V_165_A&R_166_A) with a type-II integrase mutation, and NL4.3(Ѱ: Δ(105-278)&Δ(301-332)) featuring an edited Ѱ packaging signal. Notably, virions from NL4.3(Ѱ: Δ(105-278)&Δ(301-332)) displayed a mixed population, comprising immature virions, empty cores, and cores with detectable internal density. Conversely, virions derived from NL4.3(IN: V_165_A&R_166_A) exhibited a type II integrase mutant phenotype characterized by empty cores and RNP density localized around the cores, consistent with previous studies. In contrast, virions released from pNL4-3(RT:D_185_A&D_186_A) displayed mature cores containing detectable RNP density. We suggest that the purification methods developed in this study can significantly facilitate the characterization of enveloped viruses.

## 1. Introduction

Human immunodeficiency virus (HIV) is a complex retrovirus which replicates in CD4+ T cells and macrophages, destroying the ability of the body to fight other infections and causing acquired immunodeficiency syndrome (AIDS). There are two types of HIV: HIV-1 originated from the SIV of the chimpanzee and has spread globally ^1,2^, while HIV-2 originated from the SIV of the sooty mangabey and is relatively limited to the African continent ^2,3^. Historically, purification protocols originally developed for Rous sarcoma virus (RSV) and Avian myeloblastosis virus (AMV) have been applied to purify all retroviruses, including HIV-1 and HIV-2 due to their similar density of 1.16 g/ml and ∼150 nm in diameter particle sizes ^4^. The most common purification methods are as follows: virus is first collected by centrifugation of the growth medium over a sucrose or OptiPrep solution cushion, where OptiPrep is a 60/40 solution of Iodixanol. The resulting pellet is redissolved and the particles separated over a sucrose density gradient ^4,5^, OptiPrep velocity gradient^6^, OptiPrep density gradient^7^, or OptiPrep step gradient^8^. After extraction from the gradients, virus is sedimented again and resuspended in the final buffer suitable for further analysis. Due to their pelleting and resuspension steps, these methods are ideal for purification of virions at the very high densities required for cryo electron microscopy. Retroviral purification has also been developed specifically for Lentiviral vectors^9^, and since lentiviral preps do not require the degree of concentration required for cryo electron microscopy, lentiviral vectors are commonly purified using chromatography^10^.

The primary motivation for our study was to update a purification protocol for HIV virions to capture all virions released by producer cells with minimal structural damage by calculating and minimizing the forces encountered by virions during purification. Cryo-electron tomography and single particle electron microscopy offer powerful techniques to visualize the protein-protein and protein-lipid contacts within HIV virions ^5,11–15^. The goal is to purify a majority of the virions released from producer cells so all phenotypes are captured in analysis irrespective of the variation in their structural integrity.

HIV-1 virion assembly initiates on the plasma membrane through interactions between specialized lipids, specifically PiP_2_ ^16–18^, HIV Gag proteins, and the Ѱ packaging signal of the viral genomic RNA (gRNA) ^19–25^. During its assembly, the nascent virion incorporates HIV gp40/gp120 trimers, Gag-Pol, and a number of HIV accessory proteins ^26,27^.The released virions are immature, with a distinct lattice of Gag molecules underpinning the inner leaflet of the virions^11,12^. HIV maturation is catalyzed by the active protease dimer^28^, and maturation results in the formation of a conical core made from HIV Gag capsid proteins (CA), encompassing two copies of HIV gRNA bound to Gag nucleocapsid (NC) along with the reverse transcriptase and integrase enzymes (RNP)^2,13,14^. After fusion of the viral membrane with the next host, the integrity of the mature core is essential for delivery of the HIV RNP to the nucleus of the next host^29,30^, allowing for integration of the viral DNA into the genome of the target cell. The molecular mechanism of HIV maturation is intricate and therefore offers a good target for development of next generation antivirals ^27,31^.

A major enzyme within HIV virions is the reverse transcriptase, which functions as a heterodimer, which forms after processing of Gag-Pol and release of the reverse transcriptase monomers within the lumen of HIV virions. The reverse transcriptase is essential for transcribing a single DNA molecule from the two gRNA templates embedded in each virion^2^. The _183_YMDD_186_ motif is conserved and located at the palm domain of HIV reverse transcriptase. Mutagenesis of residues 185 and 186 from aspartate to alanine abolishes RT catalytic activity and infectivity of released virions^32^. In vitro analysis of the virions with this inactivating mutation, D_185_A and D_186_A, suggested that the cores released from these virions were more stable^33^. A number of factors are involved in tuning the stability of the cores and capsid stability and reverse transcription are suggested to be balanced to minimize the innate immune sensing of HIV ^34^. Atomic force microscopy measurements also suggest stages of core stiffness spikes linked to reverse transcription ^35^. To our knowledge, there are no cryo-electron tomography characterizations of the cores generated by virions with the D_185_A and D_186_A mutation in reverse transcriptase. In contrast, the mutations within the catalytic domain of integrase have been well characterized. Specifically, mutations within the catalytic domain of integrase V_165_A and R_166_A, which result in inactivation of integrase, also create a phenotypic mislocalization of RNP outside of the mature cores within HIV virions. This phenotype which can be generated by direct mutagenesis or through action of inhibitors is defined as Type-II integrase phenotype ^2,36,37^.

Interaction between HIV genomic RNA (gRNA) Ѱ packaging signal and the nucleocapsid (NC) domain of Gag facilitate packaging of the gRNA into budding virions ^38–44^. The NC domain of Gag also has a strong affinity for cellular RNA. The absence of Ѱ packaging signal within the genome significantly reduces the amount of gRNA incorporated into HIV virions in favor of more cellular mRNAs^45,46^. Within the immature virions, the gRNA primarily binds to the NC domain of Gag^44^ and serves as a structural component of the virion^46^, after initiation of maturation, the gRNA also binds integrase, an interaction which is essential for proper maturation ^47^; however, there is no evidence that integrase would preferentially bind a specific section of the viral gRNA. There are three sections within the signal that interact with Gag, two of them are within nucleotide 105-278 and one is within 301-332, deletion of these two sections from the gRNA(ΔѰ: Δ(105 − 278)&Δ(301 − 332)) abolishes Gag binding to the packaging signal but does not affect the expression of Gag from the underlying gRNA^44^. To our knowledge, there are no cryo-electron tomography characterizations of the mature cores generated by NL4.3(ΔѰ: Δ(105 − 278)&Δ(301 − 332)) virions.

In our study we have used the pNL4.3 backbone, which encodes a genome derived from multiple wild type circulating group M HIV-1 viruses ^48^. Transfection of the pNL4.3 backbone DNA into mammalian cells, specifically HEK293 cells results in release of infectious HIV-1 virions ^49^. We present a quantitative method for HIV purification which allows the harvesting of 1,000 to 5,000 intact HIV virions from each HEK293 cell. We utilize cryo-electron tomography to compare virions produced by expression of pNL4-3(RT: D_185_A & D_186_A)^32^, NL4.3(IN: V_165_A & R_166_A)^50^ and NL4.3(ΔѰ: Δ(105 − 278)&Δ(301 − 332))^44^ in HEK293 cells.

## 2. Materials and Methods

### Cell plating and transfections

Human embryonic kidney HEK293 cells were cultured in T-25 flasks using TrypLE Express Enzyme (Gibco) and Dulbecco’s Modified Eagle Medium (DMEM) supplemented with 4 mM L-Glutamine, 4.5 g/L Glucose, sodium pyruvate (Cytiva), and 10% fetal bovine serum (Gibco). Cells were incubated at 37°C in a humidified atmosphere containing 95% air and 5% CO2 and passaged every other day or upon reaching confluency. Once the HEK293 cells reached 70-90% confluency, they were seeded into 10 cm culture dishes at 9 mL of medium per dish and incubated for 24 hours prior to transfection. At approximately 60% confluency, cells were transfected with the designated plasmid. Unless otherwise noted, transient transfections were performed using Lipofectamine 2000 reagent (Life Technology). For each transfection, 20 µg of plasmid DNA and 40 µL of Lipofectamine 2000 were separately diluted in 300 µL of Opti-MEM (Gibco) and incubated at room temperature for 5 minutes. The two solutions were then combined and allowed to incubate for an additional 20 minutes at room temperature to form DNA-Lipofectamine 2000 complexes. These complexes were added dropwise to each culture dish containing cells at 60% confluency. Cells were then incubated at 37°C for 48 hours before harvesting.

### pNL4.3 plasmid preps and mutagenesis

Cloning and plasmids: Proviral clone pNL4.3 was obtained through the NIH HIV Reagent Program, Division of AIDS, NIAID, NIH: (HIV-1), Strain NL4-3 Infectious Molecular Clone (pNL4-3), ARP-114, contributed by Dr. M. Martin. Mutagenesis resulting in pNL4-3(RT: D_185_A & D_186_A)^32^, NL4.3(IN: V_165_A & R_166_A)^50^ and NL4.3(Δ: Δ(105 − 278)&Δ(301 − 332))^44^ in the HIV-1 proviral clone pNL4-3 were performed by GenScript. For use in electron microscopy, an additional mutation was added to inhibit gp160 proteolysis, specifically NL4.3:Env(_506_SEKS_509_)^51^ to ensure biosafety during cryo-electron tomography.

DNA amplification and purification for transfection: 4ug of lyophilized plasmid was resuspended in 10uL of DNase/RNase-free distilled water and stored at –20°C. 1ul (.4ug) of each plasmid was added to 10uL of Invitrogen MAXEfficiency chemically competent DH5α cells. Plasmid and cells were incubated on ice for 30 min and then heat shocked at 42°C for 40 seconds. This was followed by a 2 minute recovery on ice. 250uL of room temperature LB was added to each tube and then plated on LB-agar plates with 0.01% ampicillin. Plates were inverted and incubated overnight at 37°C. A single colony was used to inoculate 5mL of LB with 10 ug/mL of ampicillin and incubated for 8 hours at 37°C while shaking at 225 rpm. 1mL of this starter culture was inoculated into 500mL liquid LB with 20 ug/mL of ampicillin and cultured overnight in a shaker (225 rpm) at 37°C. DNA for transfections was purified per manufacturer’s protocol, using the ThermoScientic GeneJET Plasmid Maxiprep Kit, and eluted in 1mL of elution buffer

### Preparation of OptiPrep™ – Iodixanol step gradients

Two solutions (A & B) are prepared to make the step gradient. Solution A is made from 30mL of 100mM Hepes and 70mL Phosphate Buffered Saline (PBS). Solution B consists of 10mL of 100mM Hepes and 90mL PBS. Both of these solutions are then filtered through 0.22μm filters (CellTreat Product Code: 228747) for sterilization. Then, 10mL of Solution A is mixed with 20mL of OptiPrep™ Density Gradient Medium from Sigma Aldrich. This creates a solution of 40% OptiPrep™ and will be used as the bottom step in the gradient. For the top step, mix 7.5mL of the 40% OptiPrep™ solution into 12.5mL of Solution B to make a 15% OptiPrep™ solution.

### Preparation of 10X gold 20 nm Gold beads

For use with cryo-electron tomography, functionalized gold nanoparticles must be added to the resuspended pellet at the end of the purification process. 1mL of 10 nm BSA Gold Tracers are concentrated by spinning in a 100kD Millipore Amicon Ultra, for five minutes at 5000 rpm. Then, 1mL of STE Buffer (20mM TRIS-HCl, 100mM NaCl, 1mM EDTA, pH: 7.4) is added and spun again for 3min. The desired final concentration is 10-fold over stock.

### Freezing of virions on EM Grids

Immediately following each VLP harvest, a one to one volume fraction of concentrated 10 nm gold fiducials were added to the resuspension. 3.5 uL of the resulting sample were added to glow discharged Quantifoil Ultrathin carbon R2/1 200 mesh EM Grids (Ted Pella, Redding, CA, USA; Quantifoil Micro Tools GmbH, Großlöbichau, Germany). Following a single 2 second blot and a 1 minute wait, samples were plunge frozen in a liquid ethane/propane mix, using an FEI Vitrobot Mark IV at 4℃ and 90-100% humidity.

### Cryo-electron tomography of viral particles embedded in vitreous ice

Next, cryo-specimens were imaged in a Titan Krios G3i transmission electron microscope (ThermoFisher) equipped with a Gatan BioQuantum K3 energy filter and direct electron detector. Tilt series from −60° to +60° were recorded at 3° steps over holes in the carbon film via SerialEM software. The microscope was operated at 300 kV with images having a pixel size corresponding to 1.407 Å at the specimen. Slit width of the energy filter was 20-30 eV. Tilt series were recorded at a target defocus of –5 µm. Total electron dose at the specimen was 100–125 electrons per square Å.

### Data analysis and tomogram reconstruction

To generate tilt series stacks from the raw micrographs, 4-10 frames per tilt angle were automatically aligned using SerialEM 4.1.0. The resulting 41 frame tilt series were aligned using 10-20 gold fiducials per tilt series in IMOD (University of Colorado). Using IMOD, each tilt series was then fitted for CTF correction, using Ctfplotter, and dose filtered at 80% of standard. Tomograms were then 3D CTF corrected and reconstructed in 15 nm slabs for final reconstructions of 1000-1800 slices or ∼140-250 nm total thickness.

### Measurement of Virion Retention on Blotted Grids

A 3.5 µL aliquot of a single-cycle pseudotyped HIV-1 sample (pBR4-3IeGEnv +Vsv-g) was applied to cryo-EM grids. The grids were blotted using a Vitrobot under the same conditions described for viral-like particles (VLPs) and immediately submerged in 125 µL of supplemented DMEM within a 96-well plate. Unblotted grids, which received the same volume of the sample, served as controls. Serial dilutions of the initial volume were prepared, and the media was transferred to plates containing confluent TZM-bl cells. The cells were incubated for 48 hours and subsequently stained with X-gal staining solution to quantify infectious virus particles remaining on the grids after blotting. Infectious virus titers were calculated by manually counting blue foci in the wells, then multiplying the counts by the reciprocal of the dilution factor, as previously described ^52^.

## 3. Results

### Calculating an efficient purification protocol for HIV virions

The population of HIV-1 virions produced in this study likely range in durability. To maximize virion survival in this purification protocol, our goal was to remove the virions from the supernatant with minimal applied forces. For these reasons, we chose to simulate centrifugation steps to optimize purification conditions. The simulation models are presented for two condition sets. The first set conditions for an SW41 rotor with a 13 ml centrifuge tube, which can easily accommodate the 9 ml supernatant from each 10 cm plate and the two layers of Optiprep. These densities were set at 15% and 40% Optiprep.

Purification parameters for centrifugation speed, centrifugation time, and appropriate media viscosity were tuned using a custom matlab script which simulated concentration and apparent force on the virions; the script is deposited online at (https://github.com/saveez/SaffarianLab). Virion concentration was modeled at discretized centrifugal radii, and changes to those concentrations was simulated over time using a finite difference method. At a given radius, the instantaneous particle velocity was estimated using Stoke’s law, 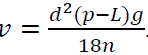. This velocity depends on the density and viscosity of the medium at that radius and the centrifugation speed. To estimate the force exerted on the virions at a particular radius, we combined this velocity with the corresponding instantaneous viscosity using the equation, 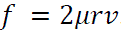. To simulate virion concentration changes over time, we calculated the instantaneous Diffusion Coefficient of the particles using the Stokes-Einstein equation, 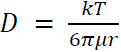, and iterated those concentration changes with the finite difference method.

Using our simulation, we show that running the centrifuge at 30000 RPM, which has an average G force of 154,000g, will collect the virions at the 15% to 40% OptiPrep solution interface within a half hour, as seen in Figure 1. The apparent force experienced by the particles, which is a function of centrifugal radius and medium viscosity, is also plotted in Figure 1. For convenience, distances are reported from the top of the buffer in the centrifuge tube at 0 and a maximum at the bottom of the tube; however in the actual calculations, distances used are distances to the center of the rotor. Maximum forces applied to particles are predicted to hit maximum immediately before they cross the gradient boundary into 15% Opti which would peak at ∼3 fN for HIV virions with a density of 1.16 and diameter of 140 nm. Once virions reach the interface between 15% and 40% Optiprep, the applied forces on the virions are zero.

**Figure 1.**
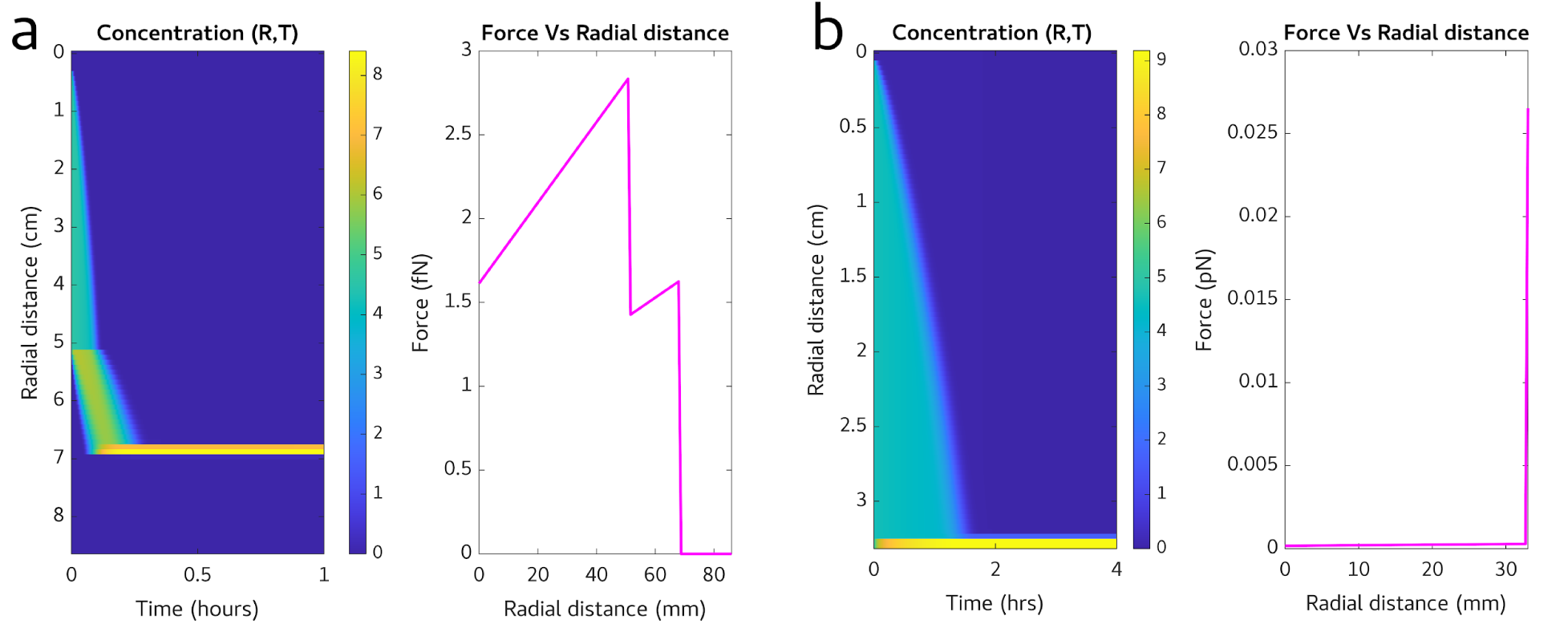
a) Simulated concentration vs time and force vs radial distance data for VLPs spun at 154,000 g over a 15%-40% OptiPrep™ step gradient. b) Simulated concentration vs time and force vs radial distance data for VLPs pelleted at 20,000 g.

After extraction of the virion band, virions are embedded in the Optiprep buffer which matches their density. By diluting the virions in PBS, one can adjust the density of the surrounding and allow for additional centrifugation to pellet the virions. We hypothesized that virion damage occurs due to the total force applied to the virion multiplied by the time the virions spend under this force. If one just dilutes the virions into a large volume, the required centrifuge speed and time for pelletting will increase, and therefore, the pelleted virions will have a higher likelihood to get damaged. As shown in Figure 1 column b, the forces applied to virions in the pellet can exceed 10 times the force experienced in the first round of centrifugation, which is a substantial amount of force. To minimize this pelleting force, we used a 3X dilution of the virion band in PBS as our working solution. This dilution allows the collected viral band and dilution to be accommodated in a 1.5ml centrifuge tube and, as shown by the calculations in Figure 1, allows for pelleting of the virions within 2 hrs at 20,000 g. The forces applied to virions at the bottom of the tube were calculated at below 0.03 pN, according to the force equation 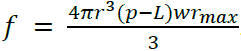. To minimizevirion damage as the particles pellet against the rigid centrifuge tube, we limited pelleting time to 2 hours, as is sufficient time for concentration according to calculations in Figure 1 b.

### Developing the experimental method for virion purification

Based on the calculation presented above, we developed the below protocol for purification as shown in Figure 2.

**Figure 2.**
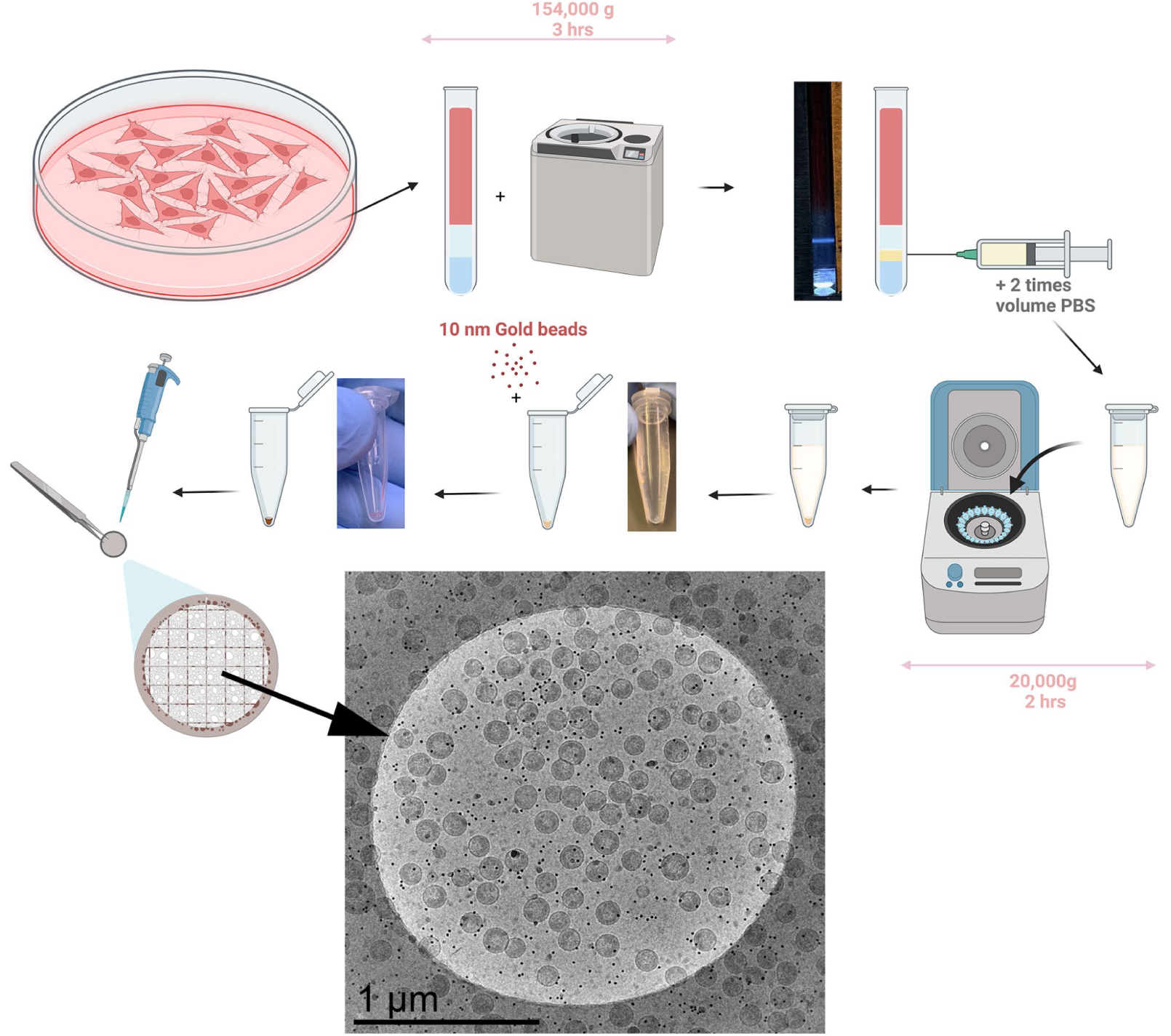
Schematic illustration of HIV VLP purification method from one plate of HEK293 cells. Images showing real samples are provided after the step-gradient centrifugation step, the pellet centrifugation step, and after adding the gold nano-particles. The final image shows a 2 um diameter circle from the final cryo-em grid.

At 48 hours post-transfection, the cells and media are removed from their plates, placed in individual 15mL tubes, and spun down at 5,000rpm for 10 minutes at 4° Celsius to pellet any cell debris. Then the supernatant is filtered through 0.22μm filters and into Ultra-Clear™ Centrifuge Tubes from Beckman Coulter (Size:14×89mm). Using a 10mL syringe with a long stainless steel loading needle, 2mL of 15% Opti Solution is very carefully added to the bottom of each centrifuge tube to create the first layer of the step gradient. The syringe and loading needle are cleaned with DI water and the previous step is repeated with the 40% Opti Solution to create the second (lower) layer of the gradient. The tubes are then balanced and spun at 30,000rpm (154,000g) for three hours at 4° Celcius.

After the spin is done, the tubes are removed and illuminated from the bottom with a custom in-house tube holder to check the quality of the virion band, for details see Figure S1. The band should be clearly visible against a dark backdrop and should have reasonably discrete edges. 0.5mL of the virion band is then slowly removed by puncturing the side of the centrifuge tube with a 20G hypodermic needle and syringe. If there are multiple tubes with the same experimental conditions, the extracted bands may be combined at this time for a higher final yield. The resulting volume of extracted virion band is then tripled by addition of PBS and subsequently spun to pellet virions. As shown in Figure 2, virions resulting from one 10cm plate will be harvested in 0.5ml, which after adding 1 ml of PBS can be spun at 20,000g for 2 hours at 4° Celcius. After this spin, the tubes are decanted into bleach and carefully dried, taking care to not dry the pellet which should be visible as shown in Figure 2. Depending on the final desired concentration, an amount of STE Buffer (20mM TRIS-HCl, 100mM NaCl, 1mM EDTA, pH: 7.4) will be introduced to the pellet. This will have to be pipetted up and down to carefully resuspend the pellet. Once the pellet has been successfully resuspended, a one to one volume of 10X concentrated gold beads in STE buffer is added before preparation of the EM grids. Figure 2, shows the purification of virions from HEK293 cells transfected with pNL4-3(D_25_N)(ΔENV) which has an inactivating D_25_N inactivating protease^53^ and a frameshifting mutation which compromises the gp160 translation. Virions shown in the figure are from one, 10cm plate prepared according to protocol shown in the figure.

### Measuring the virion yield in the purification protocol

To measure virion yield from our purification protocol, virion concentrations were extrapolated from tomograms and single capture Cryo-EM images of larger grid areas. Figure 2 shows a 1um radius circle of our Cryo-EM grid. ∼120 VLPs per 1um radius mesh circle are visible across the sample. Tomograms show an ice thickness of 140-200 nm. To measure the fraction of virions removed from the grid during blotting, we applied 3ul of a viral sample with known infectivity and measured the infectivity of the sample left on the grid after blotting. These measurements as shown in Figure S2 show a 97% reduction in infectivity after blotting. Assuming a 200 nm of vitreous ice coating on both sides of the grid, the calculated volume of liquid left on the grid is 99.92% of the applied sample. According to these calculations and measurements blotting concentrates virions onto the grid by a factor of ∼(1-0.97)/(1-0.9992)=30 fold. By applying this factor to our calculations we estimate that there are 10^12^/30 virions released from cells in a 10cm plate. Assuming a standard 8.8*10^6^ cells on a confluent 10 cm dish, this upper estimate would suggest a yield of 3800 VLPs per cell. Given uncertainty in the estimation of the thickness of ice on areas away from the holes, we estimate that the number of virions per cell are between 1000-5000 VLPs/Cell. In terms of a standard 3mm Cryo-EM grid, a single 10cm plate of confluent HEK293 cells yields a nearly complete single layer of virions for imaging using this method.

### Cryo-electron tomographic reconstruction of NL4.3:Env(_506_SEKS_509_) virions

Utilizing the above methods, we imaged virions derived from NL4.3:Env(_506_SEKS_509_). These virions are assembled from proviral DNA with identical sequence to NL4.3 except a mutation at the cleavage site of the gp160 (_506_SEKS_509_) which abrogates the cleavage of gp160 and therefore virions are released with intact gp160 in place of gp40/gp120 ^51^. We chose this mutation, primarily, because it lies within gp160 which is exposed on the outside of the virions, therefore it will have minimal impact on the maturation of HIV virions, while making the released non-infectious and safe to handle in Cryo-electron tomography. Figure 3 reveals three phenotypes observed in virions released from the NL4.3:Env(_506_SEKS_509_): a) Virions with an intact mature core and encapsidated density. b) Virions with irregular core and non-encapsidated density. c) Virions with an irregular core and encapsidated density. In comparison, 95% of analyzed virions (n=54) were type a with intact conical cores with encapsidated density. Less than 5% of virions were in either Type b or Type c category. These results are similar to previously observed phenotypes for mature cores observed using cryo-electron microscopy measurements^14^.

**Figure 3.**
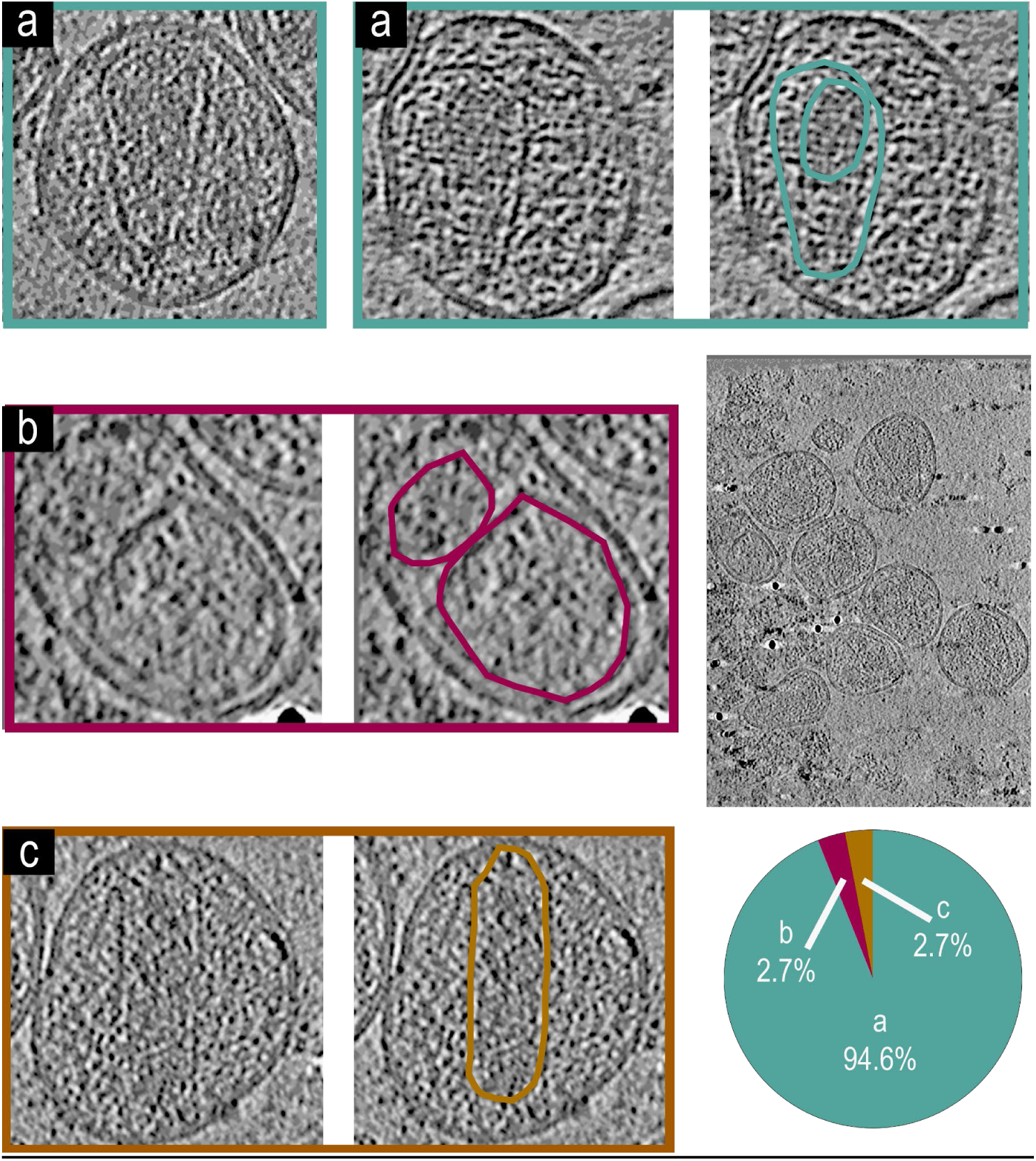
Representative tomogram slices from NL4.3:Env(_506_SEKS_509_) virions, a larger tomogram slice with multiple VLPs, and a pie chart showing phenotypic representation percentages. a) Two tomogram slices of VLPs showing a phenotype with an intact mature core and encapsidated density, both of which are highlighted in teal for the second VLP. b) A tomogram slice of a VLP showing the phenotype with an irregular core and non-encapsidated density, both of which are highlighted in magenta. c) A tomogram slice of a VLP showing the phenotype with a cylindrical core and encapsidated density, both of which are highlighted in dark yellow. (n=54) virions analyzed for the figure.

### Cryo-electron tomographic reconstruction of NL4.3(ΔѰ: Δ(105 − 278)&Δ(301 − 332)) virions

Utilizing the methods developed above, we first characterized virions generated from the NL4.3(ΔѰ: Δ(105 − 278)&Δ(301 − 332))^44^. This backbone produced 3 distinct phenotypes, two of which were previously characterized, the phenotypes are: a) Virions with an intact mature core and encapsidated density. b) Virions with irregular core and non-encapsidated density. d) immature virions. These phenotypes are represented in Figure 4. 55% of analyzed virions had the type (a) phenotype, with intact mature cores and encapsidated density, Nearly 40% of the surveyed cores (n=51) were deformed without visible density encapsidated (b), and a small portion of the remaining virions showed no signs of proteolytic cleavage and remained immature (d), These virions appear virtually identical to protease deactivated mutants^11,12^.

**Figure 4.**
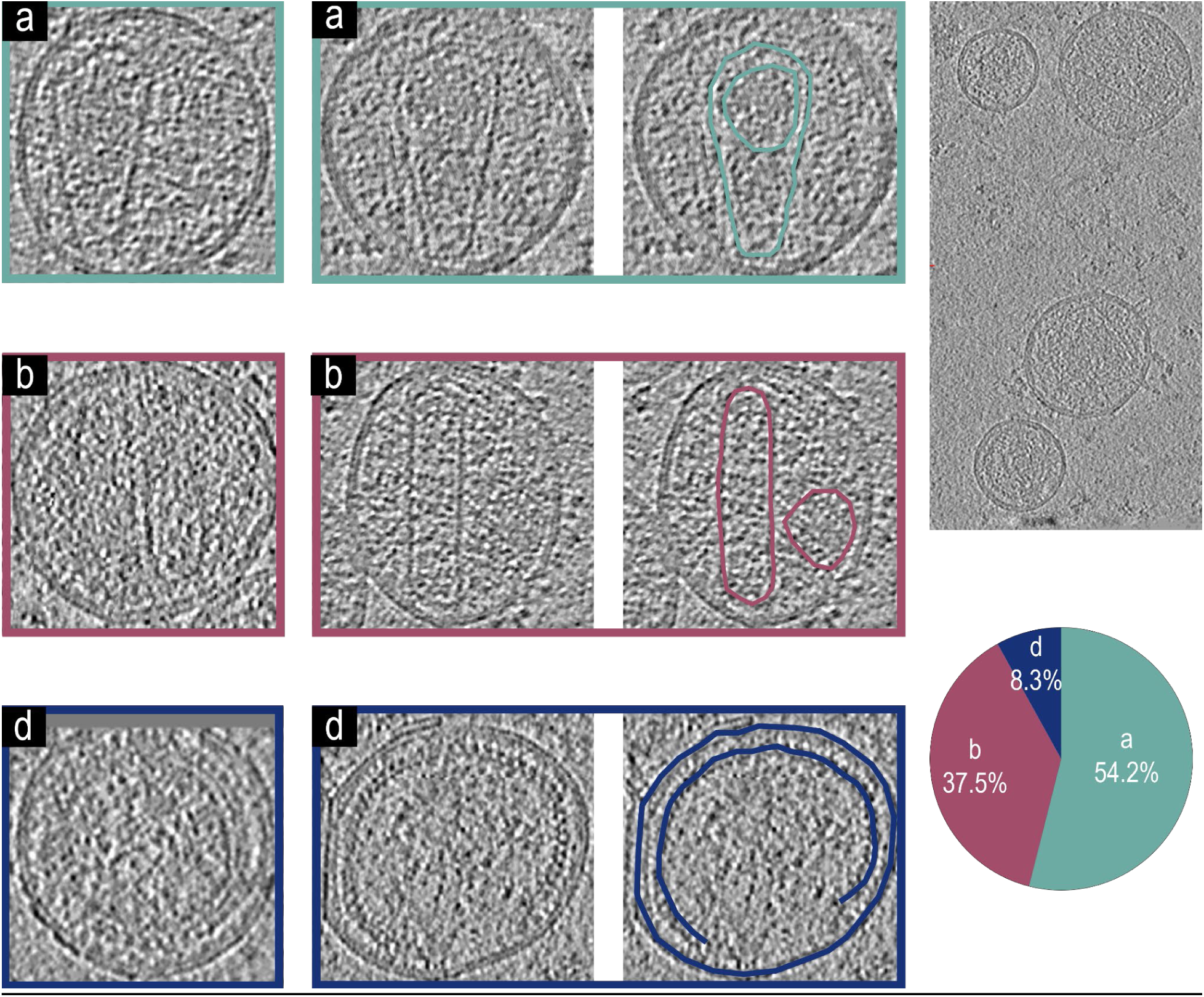
Representative tomogram slices from NL4.3(ΔѰ: Δ(105 − 278)&Δ(301 − 332)) VLPs, a larger tomogram slice with multiple VLPs, and a pie chart showing phenotypic representation percentages. a) Two tomogram slices of VLPs showing a phenotype with an intact mature core and encapsidated density, both of which are highlighted in teal for the second VLP. b) Two tomogram slices of VLPs showing a phenotype with an irregular core and non-encapsidated density, both of which are highlighted in magenta for the second VLP. d) Two tomogram slices of VLPs showing an immature phenotype without a mature core. The immature Gag lattice and viral membrane are highlighted for the second VLP in dark blue. (n=51) virions analyzed for the figure.

### Cryo-electron tomographic reconstruction of pNL4-3(RT:D_185_A & D_186_A)(Env(_506_SEKS_509_)) virions

It is known that interactions between integrase and the gRNA are essential for proper maturation of HIV, however, the effects of inactivating mutation of reverse transcriptase on the core formation have not been directly observed using cryo-electron tomographic reconstruction. Figure 5 shows two major phenotypes from the NL4-3(RT: D_185_A & D_186_A)^32^ backbone: a) Virions with an intact mature core and encapsidated density. b) Virions with irregular core and non-encapsidated density. We found that approximately 85% of observed virions (n=53) had a type (a) phenotype with mature cores encapsidating density, while 15% showing cores which were deformed and did not have encapsidated density.

**Figure 5.**
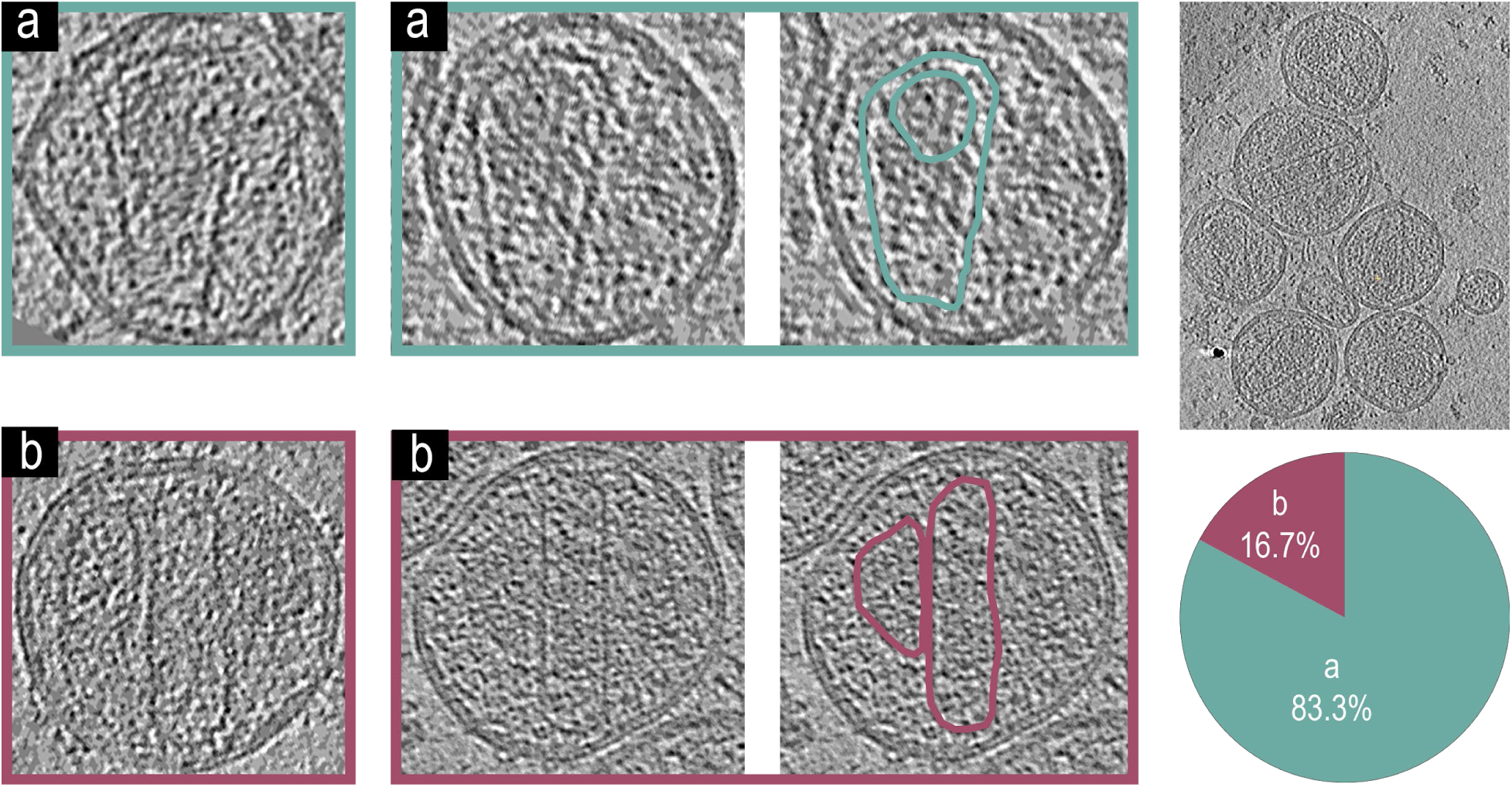
Representative tomogram slices from pNL4-3(RT:D_185_A & D_186_A)(Env(_506_SEKS_509_)) VLPs, a larger tomogram slice with multiple VLPs, and a pie chart showing phenotypic representation percentages. a) Two tomogram slices of VLPs showing a phenotype with an intact mature core and encapsidated density, both of which are highlighted in teal for the second VLP. a) Two tomogram slices of VLPs showing a phenotype with irregular cores and non-encapsidated density, both of which are highlighted in magenta for the second VLP. (n=53) virions analyzed for the figure.

### Cryo-electron tomographic reconstruction of NL4.3(IN: V_165_A & R_166_A)(Env(_506_SEKS_509_) virions

We also examined virions from the NL4.3(IN: V_165_A & R_166_A)^50^ which has a well known phenotype with mislocalization of RNP outside of the mature cores, defined as the Type-II integrase phenotype ^2,36^. Figure 6 shows the four major phenotypes from the NL4.3(IN: V_165_A & R_166_A)^50^ backbone: a) Virions with an intact mature core and encapsidated density. b) Virions with an irregular core and non-encapsidated density. d) Virions without a mature core. e) Virions with mature core and non-encapsidated density. Unlike the other backbones surveyed here, the major phenotype (58%) (n=50) of the Integrase mutant was (d) which has intact cores without encapsidated density with only 10% of virions showing type (a) phenotype with intact cores and detectable density inside. This major phenotype matches previous work identifying the Type-II integrase phenotype^2,36^.

**Figure 6.**
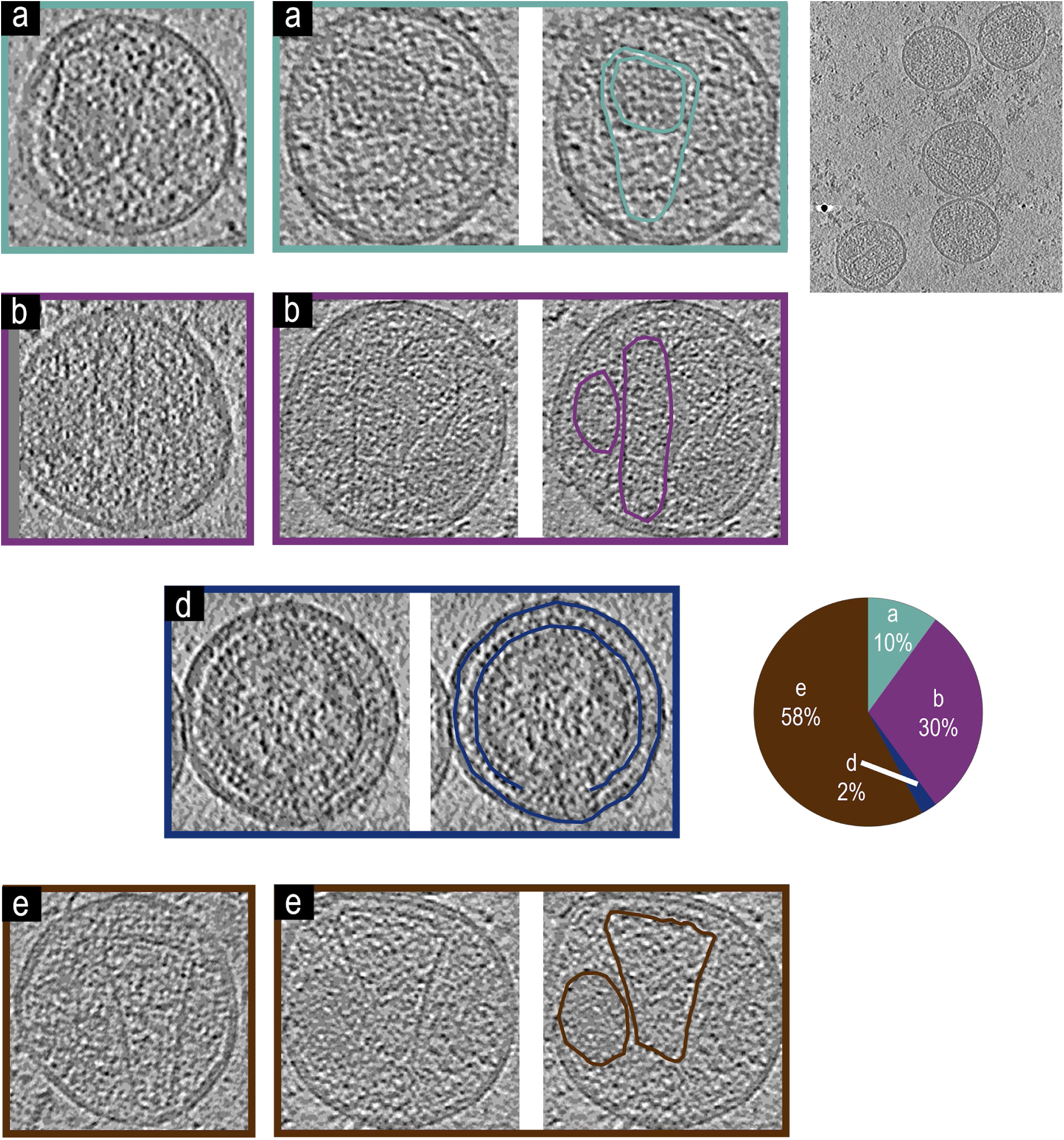
Representative tomogram slices from NL4.3(IN: V_165_A & R_166_A)(Env(_506_SEKS_509_) virions, a larger tomogram slice with multiple virions, and a pie chart showing phenotypic representation percentages. a) Two tomogram slices of VLPs showing a phenotype with an intact mature core and encapsidated density, both of which are highlighted in teal for the second virion. b) Two tomogram slices of virions showing a phenotype with an irregular core and non-encapsidated density, both of which are highlighted in magenta for the second virion. d) A tomogram slice of a virion showing a phenotype without a mature core. The immature Gag lattice and viral membrane are highlighted for the VLP in dark blue. Two tomogram slices of VLPs showing a phenotype with an intact mature core and non-encapsidated density, both of which are highlighted in brown for the second virion. (n=50) virions analyzed for the figure.

## 4. Discussion

Viral purification is an essential step in studying viruses, especially for analysis by cryo-electron tomography where highly concentrated viral samples allow efficient single particle analysis. Purifying a majority of viral particles released from cells is desirable to present a homogeneous representation of virions for structural analysis. Complex viruses like HIV are further complicated, because only 1 in 100 or 1 in 10,000 are found to be infectious ^54,55^. The current manuscript is a quantitative effort to improve the virion yield and structural integrity of released virions with the goal to improve the representation of virions in analysis independent of their structural stability. The proposed protocol is also efficient and short, allowing preparation of one cryo-EM grid of high density virions from a 10 cm dish of HEK293 cells producing HIV virions from NL4.3 backbone, within as little as 6 hrs (see figure 2).

In our estimate of virion purification we arrive at 1,000-5,000 virions released from individual HEK293 cells. It’s important to discuss the approximations we used to arrive at this number. We measure the density of virions which are captured at the end of the protocol in vitreous ice on electron microscopy grids, where we could also measure the thickness of the vitreous ice using tomographic analysis of the sample. As mentioned in the results, we also estimated how virions got concentrated during blotting by estimating the thickness of the vitreous ice remaining on the grids and comparing this with the loss of infectivity accompanying the blotting process to arrive at an approximately 30 fold concentration enhancement of virions during the blotting process, because we do not have accurate data on the thickness of the ice away from the observable areas in the grid, we had to make assumptions which resulted in our higher and lower bounds of 1,000 and 5,000.

The release of the virions is catalyzed by endosomal sorting complexes required for transport (ESCRTs)^56–65^. The released virions are immature, with a distinct lattice of Gag molecules underpinning the inner leaflet of the virions^11,12^. HIV maturation is catalyzed by the active protease dimer^28^, which needs to form by releasing the protease monomers embedded within the Pol portion of the Gag-Pol within the lumen of immature virions^2,66^. What regulates Gag-Pol auto-processing and where the Gag-Pol proteins are located within the lattice of the immature virions remains unknown. HIV maturation and protease activation coincide with release of the virions and over packaging of Gag-Pol or delay in release of the virions results in premature activation of the protease and release of non-infectious virions^67,68^.

How does the release of 5,000 virions per HEK293 cells compare with previously measured yields? Experiments with Rous Sarcoma Virus have shown that during active infection of cultures, cells can dedicate approximately 1% of their total protein synthesis to producing RSV virions^4^. Here we will derive a back of the envelope comparison to HEK293 cells. In their growth phase, HEK293 cells double every 24 hours, and therefore cells are capable of producing an equal mass of their biological material in 24 hours. HEK293 cells have a mass of 5 10^-10^g ^69^ and the mass of a single HIV virion is 10^-18^g and therefore the upper bound estimate of HIV production can not exceed 10^6^ virions per cell. We estimated harvesting 5×10^3^-1×10^3^ virions per HEK293 cell which would be approximately 0.5%-0.1% of their biomass production respectively. These numbers are in reasonable agreement with measurements of RSV production.

We used this method of purification to analyze virions released from HEK293 cells transfected with pNL4-3(RT: D_185_A & D_186_A)^32^, NL4.3(IN: V_165_A & R_166_A)^50^ and NL4.3(ΔѰ: Δ(105 −278)&Δ(301 − 332))^44^. Virions released from NL4.3(IN: V_165_A & R_166_A) have been previously analyzed and identified with type-II integrase phenotype which we have reproduced in our data. The observation that virions released from NL4.3(IN: V_165_A & R_166_A) have intact cores with RNP inside, is new, but not surprising, since to our knowledge, there is no other data suggesting that RT plays a role in core formation. The mix phenotype observed with NL4.3(ΔѰ: Δ(105 − 278)&Δ(301 − 332)) is however perplexing partly because of observation of immature virions. The virions generated for this experiment are assembled in HEK293 cells from a single proviral DNA missing only the two section of nucleotides which represent the Δ packaging signal and therefore we expect any cell that would express Gag, would also express Gag-Pol, with both Gag and Gag-Pol proteins incorporating within virions with similar ratios as observed from the parental NL4.3 backbone. If there are Gag-Pol’s incorporated within these virions, then why are some of the virions immature is perplexing. Previously it has been suggested that incorporation of Gag-Pol’s is dependent on incorporation of gRNA^70^, which can explain our observation, however in this study we do not provide any additional evidence and so this remains speculative.

## Acknowledgment

Funding: This research was funded by NIH R56 AI150474-06A1 & NIH R01AI186663 to (SS). The following reagent was obtained through the NIH HIV Reagent Program, Division of AIDS, NIAID, NIH: TZM-bl Cells, ARP-8129, contributed by Dr. John C. Kappes, Dr. Xiaoyun Wu and Tranzyme Inc.”

