## Supplemental figures and text for "Enhanced Yield and Gentle Purification of HIV for Cryo-Electron Tomography Analysis of Virion Maturation"

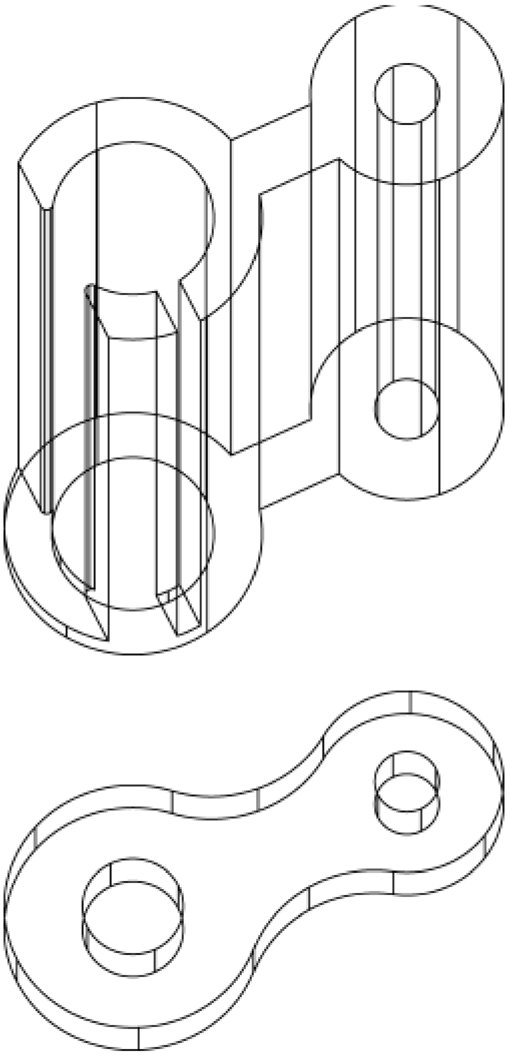

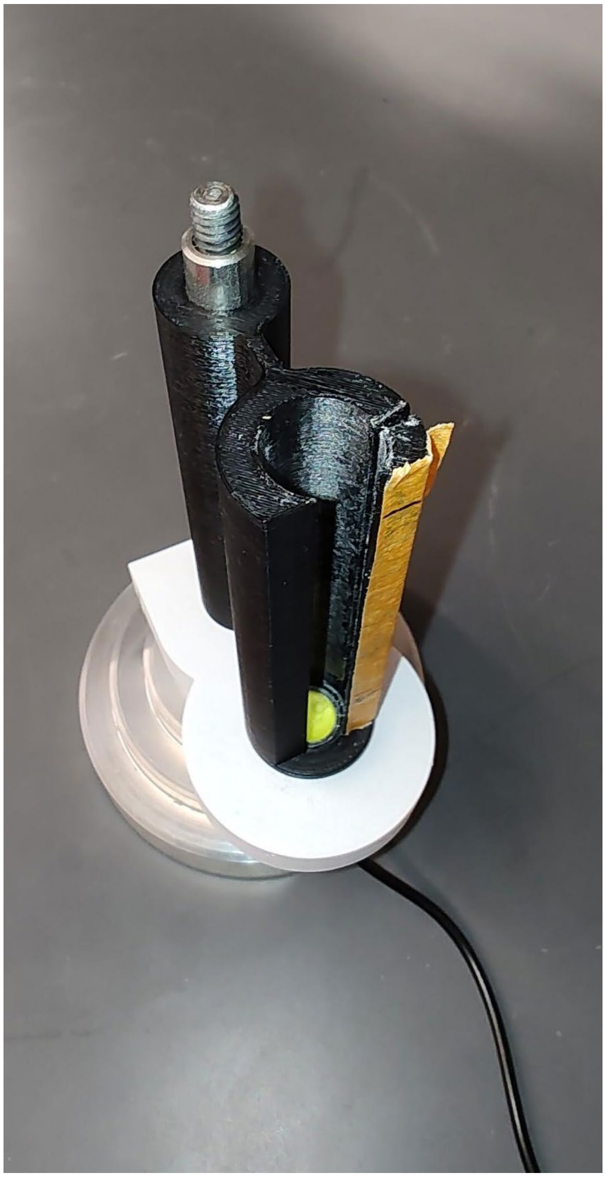

**Figure S1:** Device constructed for extraction of viral bands from the step gradients. The design allows a 13 ml clear plastic centrifuge tube to be illuminated from the bottom with an LED (The green bulb at the bottom of the chamber) as shown in the image on the right. Left shows the design of the holder from the CAD file used to construct the device.

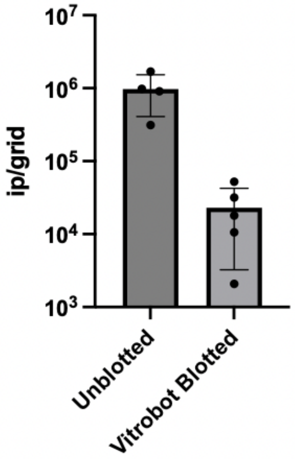


| | Unblotted | Vitrobot Blotted |
|---|---|---|
| Mean | 972656 | 22925 |
| Std. Deviation | 563133 | 19678 |
| Std. Error of Mean | 281566 | 8800 |

**Figure S2:** Measure of virions left in the grid after blotting. Number of infectious particles were quantified using blue-foci assay. 97.7% of viruses are removed by the blotting conditions described in the Methods section.
